# Inhibition of CDK4/6 Overcomes Primary Resistance to PD-1 Blockade in Malignant Mesothelioma

**DOI:** 10.1101/2021.03.04.432811

**Authors:** Hee-Jin Jang, Cynthia Y. Truong, Eric M. Lo, Hudson M. Holmes, Daniela Ramos, Maheshwari Ramineni, Ju-Seog Lee, Daniel Y. Wang, Massimo Pietropaolo, R Taylor Ripley, Bryan M. Burt, Hyun-Sung Lee

## Abstract

**Background:** Despite the profound number of malignant pleural mesothelioma (MPM) patients now treated with PD-1 blockade, insight into the underpinnings of rational therapeutic strategies to treat resistance to checkpoint immunotherapy remain unrealized. Our objective was to develop a novel therapeutic approach to overcome primary resistance to PD-1 blockade in MPM.

**Methods:** We generated a transcriptome signature of resistance to PD-1 blockade in MPM patients treated with nivolumab (4 responders and 4 non-responders). We used the TCGA MPM cohort (N=73) to determine what genomic alterations were associated with the resistance signature. We tested whether regulation of identified molecules could overcome resistance to PD-1 blockade in an immunocompetent mouse malignant mesothelioma model.

**Results:** Immunogenomic analysis by applying our anti-PD-1 resistance signature to the TCGA cohort revealed that deletion of *CDKN2A* was highly associated with primary resistance to PD-1 blockade. Under the hypothesis that resistance to PD-1 blockade can be overcome by *CDK4/6* inhibition, we tested whether *CDK4/6* inhibitors could overcome resistance to PD-1 blockade in subcutaneous tumors derived from *Cdkn2a(−/−)* AB1 malignant mesothelioma cells, which were resistant to PD-1 blockade. The combination of daily oral administration of *CDK4/6* inhibitors (abemaciclib or palbociclib) and intraperitoneal anti-PD-1 treatment markedly suppressed tumor growth, compared with anti-PD-1 or *CDK4/6* inhibitor alone.

**Conclusions:** We identified a novel therapeutic target, *CDK4/6*, to overcome primary resistance to PD-1 blockade through comprehensive immunogenomic approaches. These data provide a rationale for undertaking clinical trials of *CDK4/6* inhibitors in the more than 40% of patients with MPM who demonstrate loss of *CDKN2A*.

## INTRODUCTION

Immune checkpoint inhibitors (ICIs) have dramatically changed the treatment approach to various advanced cancers, resulting in durable clinical responses in many cases.^1–4^ Programmed Cell Death 1 (PD-1) inhibitors have recently shown encouraging clinical activity with good tolerability in patients with advanced malignant pleural mesothelioma (MPM) previously treated with chemotherapy. In these single-agent trials, objective response rates, defined as complete response (CR) or partial response (PR), ranged from 15 to 21%; and rates of stable disease (SD) were 33 to 56%, indicating a favorable impact of these drugs on durable clinical benefit.^5–7^ However, similar to patients with other tumor types treated with anti-PD-1 therapy, most patients with MPM do not respond to PD-1 inhibition, highlighting an unmet need for the development of novel therapeutic approaches to augment the efficacy of immune checkpoint treatment.

Anti-PD-1 monoclonal antibody blocks the interaction between PD-1 on activated T cells and its ligands, which is thought to reinvigorate T cells to kill cancer cells.^8,9^ However, the therapeutic efficacy of PD-1 checkpoint inhibition is limited by primary and acquired resistance.^10–13^ Resistance to ICIs may be classified into two categories: (1) primary resistance, which generally refers to patients who do not respond at all to ICIs and instead progress; and (2) acquired resistance referring to those who have an initial response to ICIs followed eventually by disease progression. The mechanisms responsible for primary resistance continue to be characterized as ineffective T cell priming, lack of tumor recognition due to defective antigen presentation, suppression via other checkpoints, tumor cell resistance to interferon, and local immunosuppressive factors in the tumor-immune microenvironment.^14^ Furthermore, there is emerging data pointing to the loss-of-function mutations in JAK1 and JAK2 from the IFN signaling pathway and beta-2 microglobulin (B2M) from the antigen presentation pathway in patients who develop acquired resistance to immune checkpoint blockade therapy.^15–18^

To date, there have been no approved therapeutic breakthroughs for thwarting resistance, and there has been a remarkably unmet need to investigate the mechanisms of resistance to ICIs. Understanding the underlying mechanisms of resistance may allow the rational design of combinatorial therapies to overcome primary resistance. Therefore, mechanism-based strategies to overcome resistance to PD-1 blockade therapy are urgently needed. Our objective was to develop a novel therapeutic approach to overcome resistance to PD-1 blockade in MPM through comprehensive immunogenomic analysis.

## MATERIAL and METHODS

### Patients and Data Acquisition

This study was performed in accordance with Institutional Review Board protocol at Baylor College of Medicine (H-43208). Informed consent was obtained for the collection of clinical data and biospecimens. We performed transcriptome mRNA arrays using the pre-immunotherapy tumor biopsies of eight MPM patients with advanced and unresectable MPM, whom we treated with nivolumab (anti–PD-1) therapy after they had progressed following treatment with a platinum-based agent and pemetrexed (**Supplementary Table 1**).^19^ Original mRNA expression data deposited in the National Center for Biotechnology Information’s (NCBI) Gene Expression Omnibus database (GSE99070) were used to generate a transcriptome-based anti-PD-1 resistance signature. To identify the association of genetic alteration with this anti-PD-1 resistance signature, we utilized mRNA sequencing data from 73 samples in The Cancer Genome Atlas (TCGA) portal (https://gdc-portal.nci.nih.gov/).^20^

Briefly, anti-PD-1 resistance signature was used to generate a support vector machine (SVM) classifier^21^ for estimating the likelihood that a particular MPM tumor belonged to the subgroup in which the anti-PD-1 resistance signature is present (anti-PD-1 resistant subgroup) or the subgroup in which the signature is absent (anti-PD-1 responsive subgroup). The robustness of the classifier was estimated by the misclassification rate determined during leave-one-out cross-validation (LOOCV) of the training set. After LOOCV, the sensitivity and specificity of the prediction models were estimated by the fraction of samples correctly predicted.

### Mouse malignant mesothelioma cell line and syngeneic mouse tumor model

AB1 mouse malignant mesothelioma (MM) cell lines, purchased from Sigma-Aldrich, were thawed and cultured at 37°C with 5% CO_2_ in RPMI 1640 (with 2mM L-Glutamine+25mM HEPES) supplemented with 5% fetal bovine serum, 100 U/mL penicillin, and 100 μg/mL streptomycin.

Six-week-old male BALB/cJ mice (Jackson Laboratory) were used under the protocol approved by the Institutional Animal Care and Use Committee, in accordance with all relevant animal-use guidelines and ethical regulations. To develop the immunocompetent mouse model of MM, 5×10^5^ tumor cells in 50 μL of the serum-free medium were inoculated subcutaneously on the right flank area, where was prepared with 70% alcohol. Tumor size was monitored three times per week. The greatest longitudinal diameter (length) and the greatest transverse diameter (width) were determined to determine by external caliper and used to calculate tumor volume. The tumor volume, based on these caliper measurements, was calculated using the modified ellipsoidal formula: tumor volume = ½(length×width^2^).

Before *in vivo* drug treatment, mice were randomly divided into 6 groups when the length of tumor of each mouse reached 5mm. For anti-PD-1 therapy, anti-mouse PD-1 Abs (10mg/kg, clone 29F.1A12, BioXCell, Cat# BE0273) or isotype control (Clone 2A3, BioXCell, Cat# BE0089) were injected intraperitoneally twice per week. For cyclin-dependent kinase (CDK) 4/6 inhibition, abemaciclib (Cat# S5716) and palbociclib (Cat# S1116) was purchased from Selleckchem. Abemacoclib (75mg/kg) and palbociclib (100mg/kg) dissolved in normal saline were administered by oral gavage daily. Mice were euthanized three weeks after the first treatment to measure tumor burdens.

Methods for Imaging mass cytometry (IMC) and time-of-flight mass cytometry (CyTOF) were described in the supplementary section (**Supplementary Table 2 and 3**).

### Statistical Analysis

Student’s t-tests or Mann-Whitney U tests were used to compare continuous variables, and Chi-square or Fisher’s exact tests were performed to analyze categorical variables. Z values were calculated by subtracting the overall average gene expression from the raw data for each gene and dividing that result by the standard deviations of all of the measured expression. All statistical analyses were performed with SPSS 27.0 (IBM Corp, Released 2020. IBM SPSS Statistics for Windows, Version27.0. Armonk, NY), Prism 5.0 (GraphPad Software, Inc, San Diego, CA), and R language and software environment version 4.0.3 (http://www.r-project.org). Statistical significance was considered at P<0.05, and all tests were two-tailed.

## RESULTS

### Development of anti-PD-1 resistance mRNA signature

This study scheme is illustrated in **Figure 1**. We first designed an anti-PD-1 resistance mRNA signature representative of resistance to PD-1 checkpoint immunotherapy by comparing mRNA expression in 4 responders (CR+PR) and 4 MPM patients with progressive disease after nivolumab treatment. Differential gene expression between responders and progressors was found in 158 differential mRNAs with a p-value less than 0.01 and above 2-fold changes, which was defined as our anti-PD-1 resistance signature (**Fig.2A** and **Supplementary Table 4**). Exploratory analysis of upstream regulators through the Ingenuity pathway analysis (http://www.ingenuity.com) revealed that the anti-PD-1 resistance gene signature was significantly associated with CDK4/6 and cyclin D1 activation in the cell cycle pathway (**Fig.2B**).

**Figure 1.**
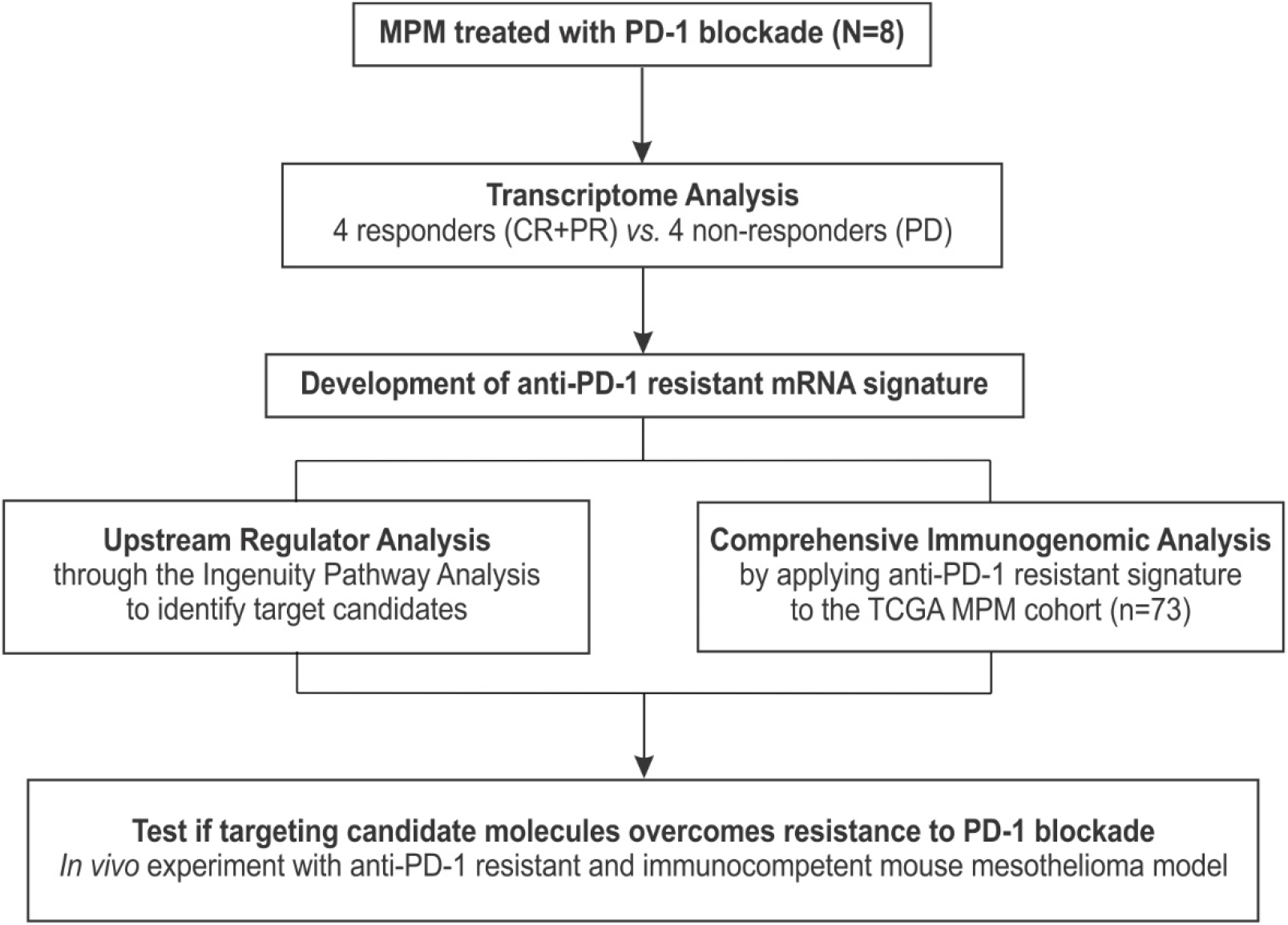
Study scheme. A transcriptome signature of resistance to PD-1 blockade was generated from malignant pleural mesothelioma (MPM) patients treated with nivolumab. To identify target candidates to overcome resistance, the upstream regulator analysis was performed. Next, the TCGA MPM cohort was used to determine what genomic alterations are associated with immune resistance signature. Finally, to test if targeting candidate molecules can increase the efficacy of PD-1 blockade, we performed *in vivo* experiment. (CR, complete response; PD, progressive disease; PD-1, programmed cell death 1; PR, partial response; TCGA, The Cancer Genome Atlas)

**Figure 2.**
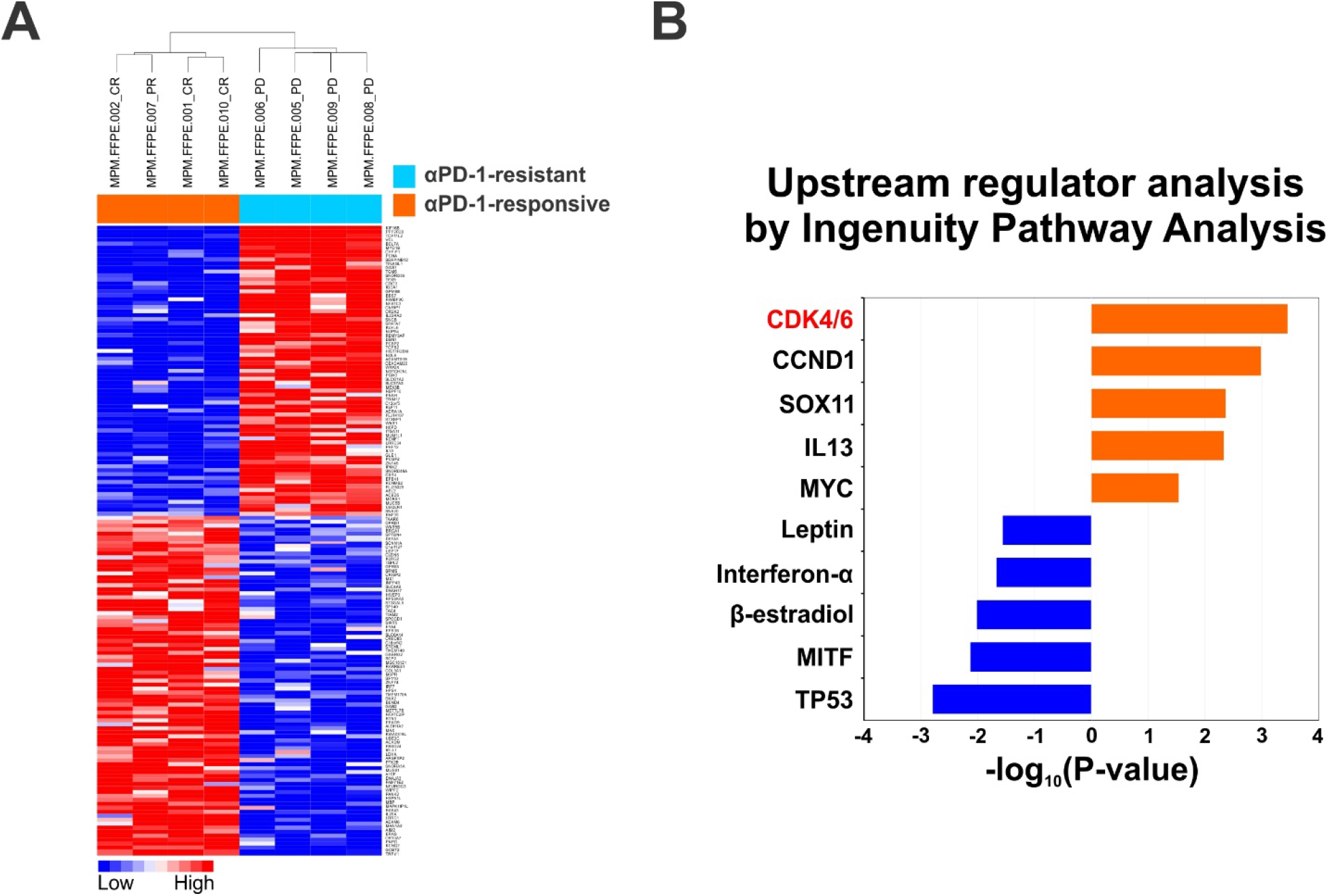
Development of anti-PD-1 resistance mRNA signature. **A**. Development of transcriptome-based anti-PD-1 resistance signature. The data are presented in a matrix format in which rows represent individual gene and columns represent individual experiment. The red and blue colors in the cells reflect relative high and low expression levels, respectively, as indicated in the scale bar (log2 transformed scale). **B**. Upstream regulator analysis through the Ingenuity pathway analysis revealed that anti-PD-1 resistant gene signature was significantly associated with Cyclin-dependent kinase 4/6 and cyclin D1 activation in cell cycle pathway.

### Resistance to PD-1 blockade is associated with CDKN2A deletion

To determine the genomic alterations associated with the anti-PD-1 resistance signature, our immune signature was applied to predict the subgroups in the TCGA MPM cohort (N=73), utilizing a previously developed model (**Fig.3A**).^19,22,23^ No clear associations of clinical features were observed between two subgroups. BAP1 mutation was observed only in the anti-PD-1 responsive subgroup (P=0.016). Furthermore, anti-PD-1 responsive tumors had abundant plasma cells and M1 macrophages, which were calculated using CIBERSORT (http://cibersort.stanford.edu/), an analytic deconvolution method of “*in silico* flow cytometry” that accurately estimates the abundances of specific leukocyte subsets using gene expression data.^24^ In contrast, MPM tumors predicted as anti-PD-1 resistant tumors had a significant deep deletion of Cyclin Dependent Kinase Inhibitor 2A (*CDKN2A*) located at 9p21.3. *CDKN2A* loss was found in 47% (N=34) of MPM patients. Among them, 74% (N=25) were classified into anti-PD-1 resistant tumors. Resting memory CD4 T cells were significantly increased in this subgroup (**Fig.3B**). Representative multiplexed images of an MPM patient with intact *CDKN2A* who had a partial response to nivolumab, showed abundant infiltration of CD8 T cells and B cells. In contrast, the other representative images of an MPM patient with *CDKN2A* loss who had progressive disease, showed abundant infiltration of tumor-associated macrophages (TAMs) in the absence of CD8 T cells (**Fig.4**).

**Figure 3.**
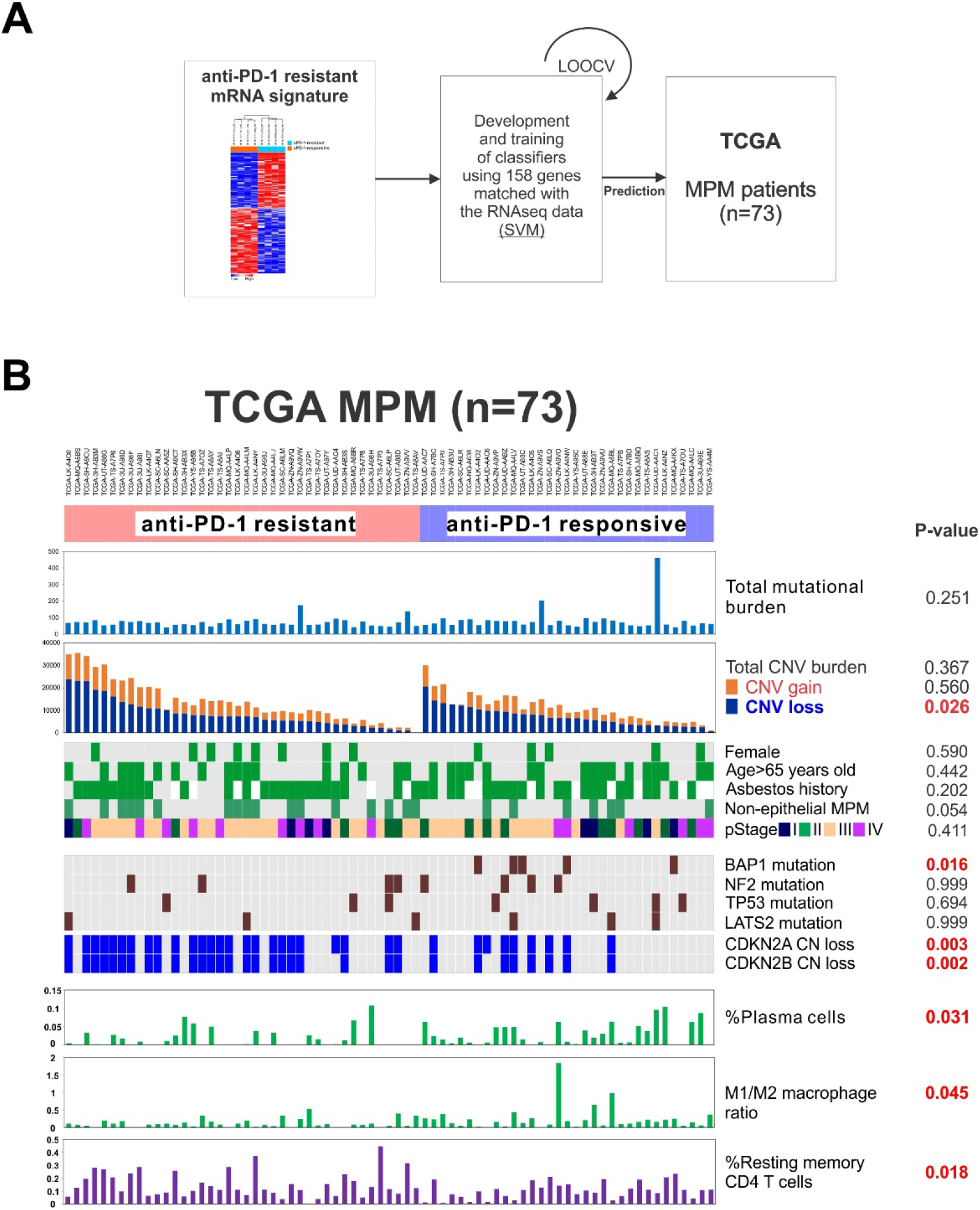
CDKN2A deletion is associated with αPD-1 resistance. **A.** The anti-PD-1 resistant mRNA signature was used to predict the subgroup in 73 TCGA MPM cohort. **B.** MPM tumors predicted as anti-PD-1 resistant tumors had a significant deletion of CDKN2A located at 9p21.3. CDKN2a loss was found in 47% of MPM patients. Among them, 74% were classified into anti-PD-1 resistant tumors. (LOOCV, leave-one-out cross validation; SVM, support vector machine; TCGA, The Cancer Genome Atlas)

**Figure 4.**
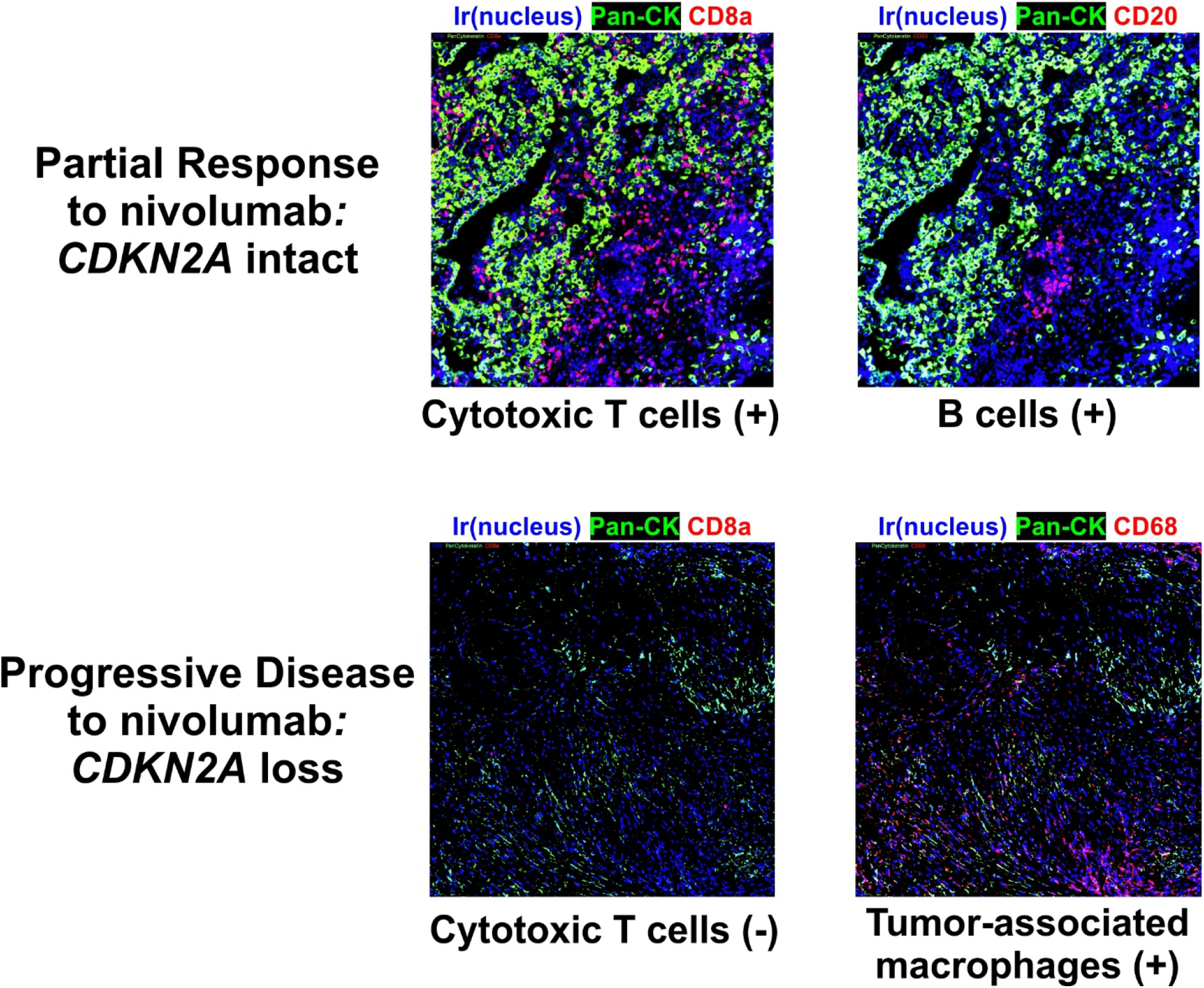
Representative images to show the association of CDKN2A status with the tumor-immune microenvironment. Representative images of an MPM patient with intact CDKN2A who had a partial response, showed abundant infiltration of CD8 T cells and B cells. In contrast, the other representative images of an MPM patient with CDKN2A loss who had progressive disease, showed abundant infiltration of TAMs with the absence of CD8 T cells. (Ir, iridium; Pan-CK, pan-cytokeratin.)

In addition, antigen presenting machinery including *HLA-A, HLA-B, HLA-C,* and *B2M* was significantly downregulated in the anti-PD-1 resistant subgroup (**Fig.5A)**. The cytolytic activity^25^ derived from the geometric mean of *GZMA* and *PRF1*, and the *IFN-γ* signaling signature (*CIITA, GBP4, GBP5, IRF1, IRF2,* and *JAK2*)^26^ were significantly lower in the anti-PD-1 resistant subgroup, compared with those of the anti-PD-1 responsive subgroup (**Fig.5B**). On the contrary, *CDK4* and *CDK6* mRNA expression was significantly increased in the anti-PD-1 resistant subgroup (**Fig.5C**), implying cell cycle activation through the depletion of *CDKN2A*.

**Figure 5.**
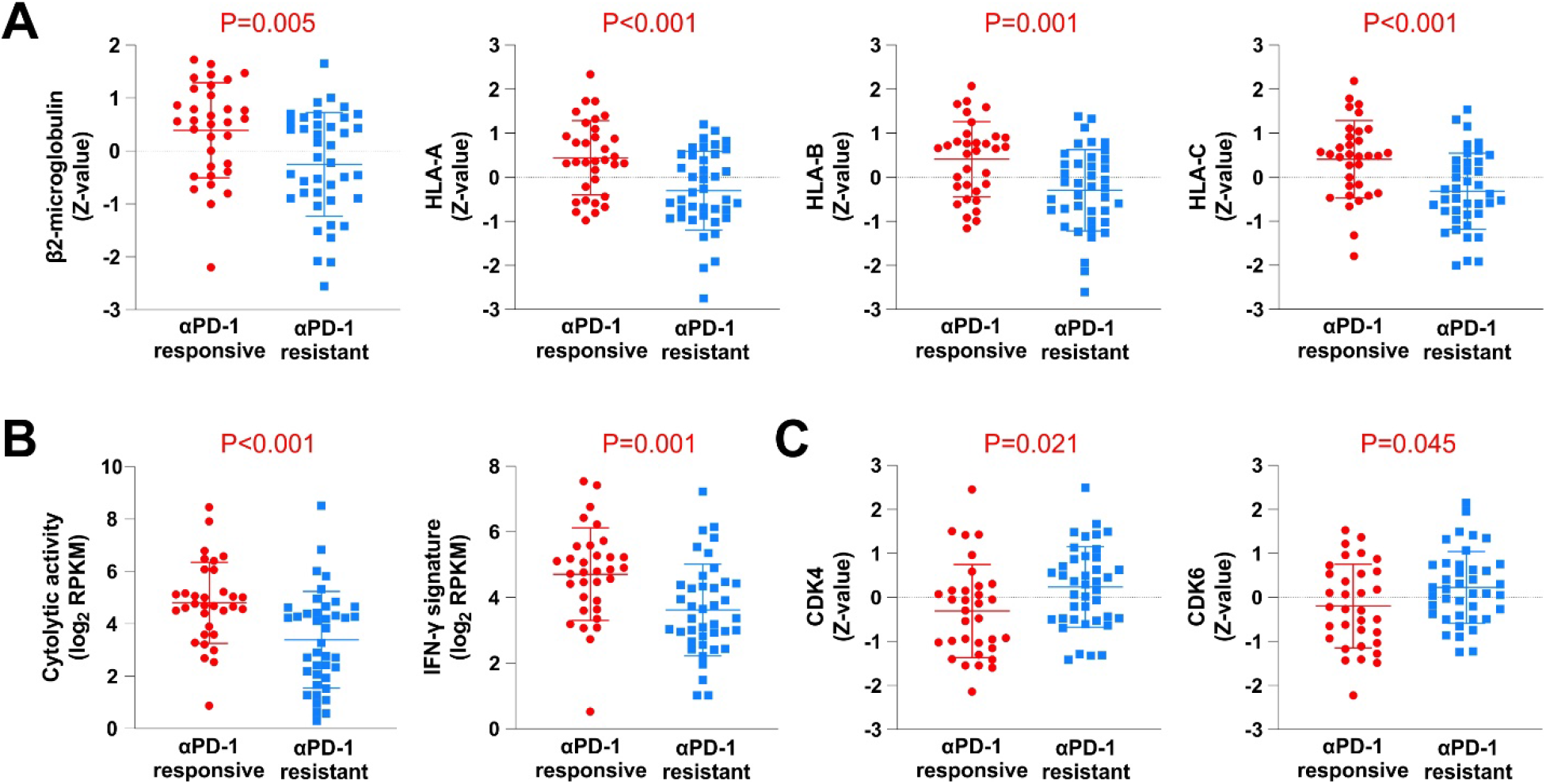
Comparison of mRNA expression related to anti-PD-1 response and resistance. **A.** The expression of B2M and MHC I molecules was significantly reduced in the anti-PD-1 resistance group. **B**. Cytolytic activity and IFN-γ signature. Subset predicted as anti-PD-1 resistance demonstrated significantly decreased cytolytic activity and the IFN-γ signaling signature. **C**. Cell cycle-related CDK4/6 expression. (RPKM, Reads Per Kilobase of transcript, per Million mapped reads.)

### CDK4/6 inhibitors overcome resistance to PD-1 checkpoint immunotherapy

*CDKN2A* is a well-known tumor suppressor that inhibits CDK4/6. The *CDKN2A-cyclin D-CDK4/6-retinoblastoma* protein pathway promotes G1-S cell-cycle transition, and this pathway is commonly dysregulated in most cancers. CDK4/6 inhibitors leading to cell-cycle arrest, senescence, and/or cell apoptosis are well described in preclinical models of various cancers, including melanoma and breast cancer.^27,28^ Increased activity of CDK4/6 is seen where *CDKN2A* is deleted.^29,30^ Therefore, we hypothesized that CDK4/6 inhibition could sensitize the tumor-immune microenvironment to PD-1 blockade. Whole exome sequencing of fifteen asbestos-induced murine MM tumor cell lines from BALB/c, CBA, and C57BL/6 mouse strains showed homozygous loss of *Cdkn2a* in 14 out of 15 tumors including AB1 MM cells, which demonstrated resistance to PD-1 blockade *in vivo* (**Fig.6A**).^31^ To test our hypothesis, we used a subcutaneous mouse malignant mesothelioma (MM) model using the AB1 cell line. Like human MPM resistant to PD-1 blockade, this mouse MM tumor contained abundant TAMs with less lymphocyte infiltration (**Fig.6B**).

**Figure 6.**
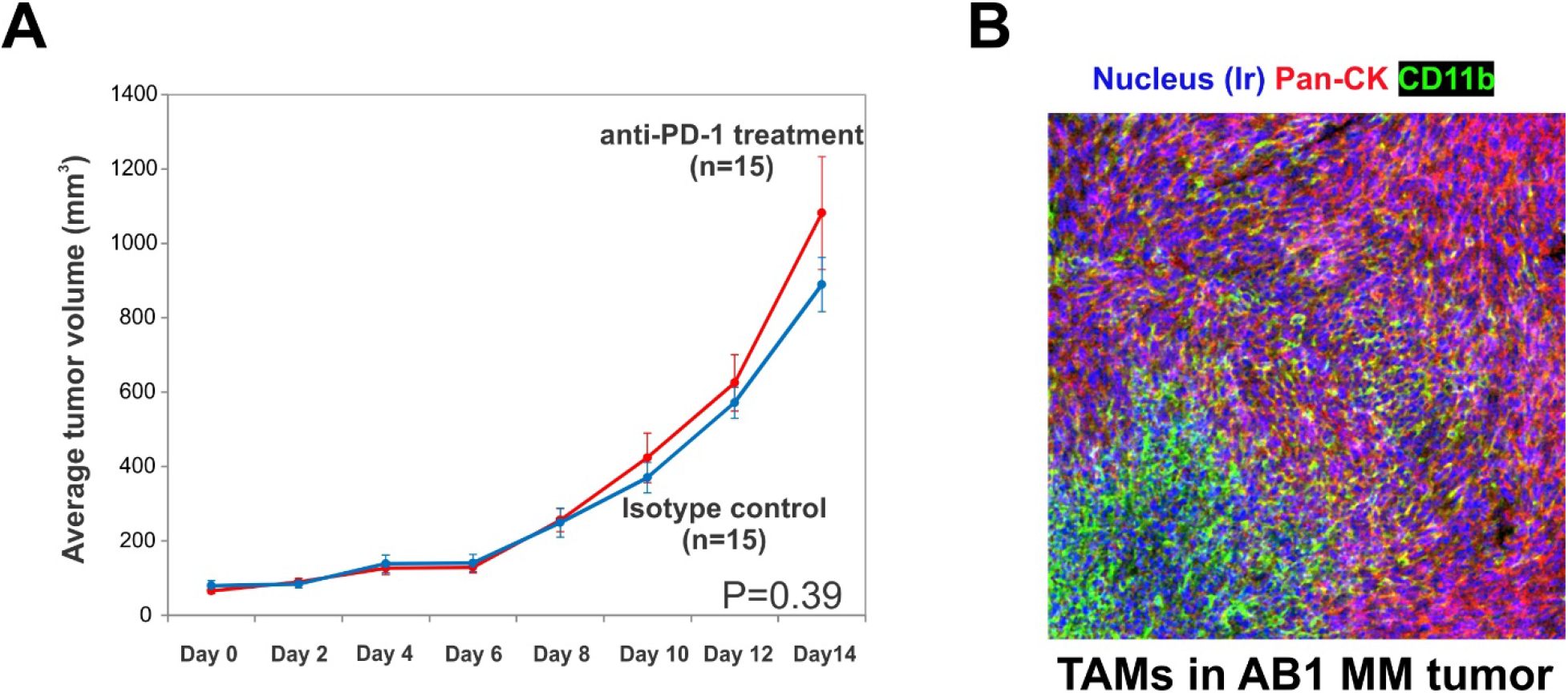
Anti-PD-1 resistant mouse malignant mesothelioma model using *Cdkn2a*(-) AB1 cell line. **A**. AB1 MM tumors were resistant to PD-1 blockade *in vivo*. **B**. Like human MPM resistance to PD-1 blockade, this mouse mesothelioma tumor contained abundant tumor-associated macrophages (TAMs) with less lymphocyte infiltration.

FDA-approved oral CDK4/6 inhibitors, palbociclib (100mg/kg) and abemaciclib (75mg/kg), were administered daily 7 days after tumor inoculation for 3 weeks, and anti-mouse PD-1 antibody (10mg/kg) was administered intraperitoneally twice per week (**Fig.7A**). CDK4/6 inhibition slowed tumor growth compared with untreated and anti-PD-1 treated groups. Abemaciclib or palbociclib single treatment showed modest effects for controlling tumor growth. A combination of CDK4/6 inhibitor and PD-1 blockade markedly suppressed tumor growth (both P<0.001)(**Fig.7B-7C**).

**Figure 7.**
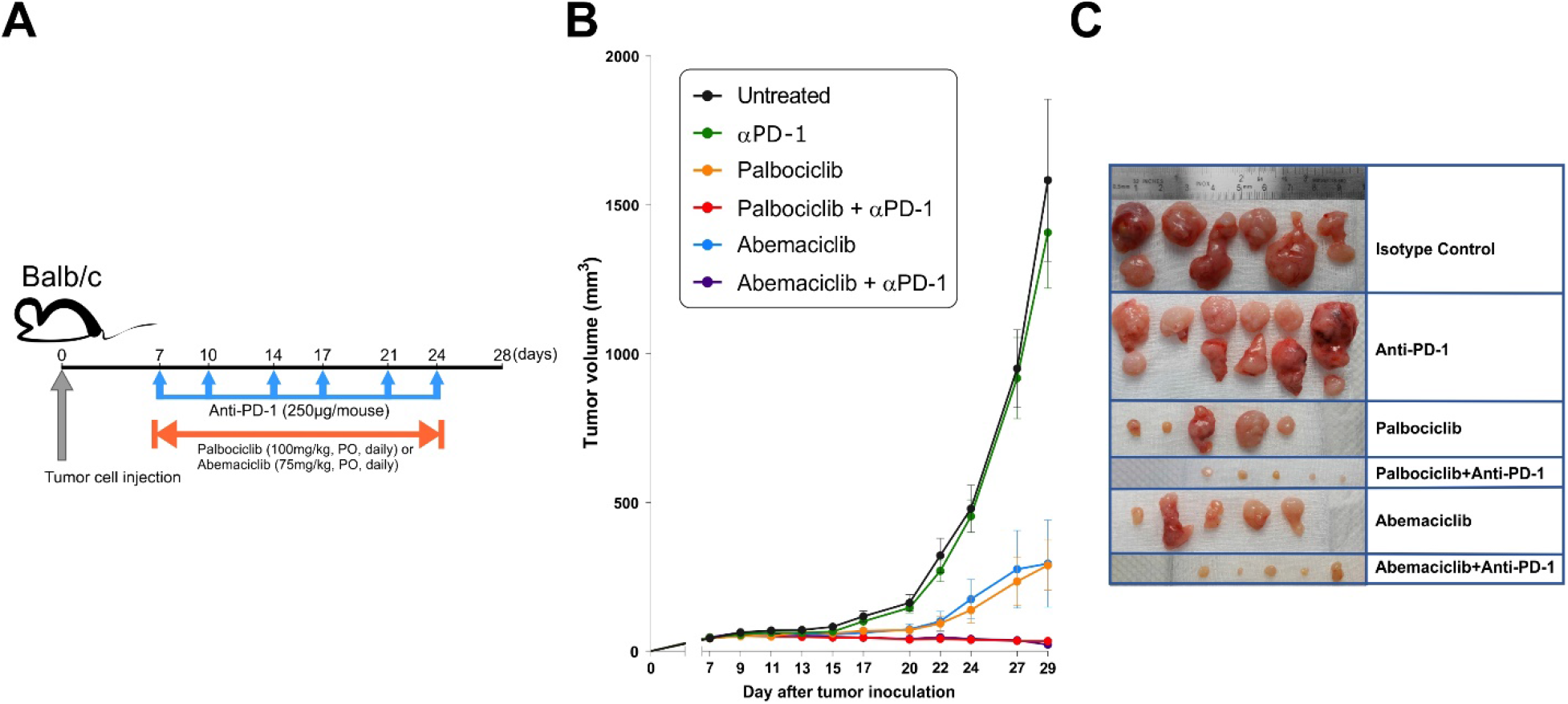
CDK4/6 inhibitors overcome resistance to PD-1 blockade. **A**. Treatment scheme. FDA-approved oral CDK4/6 inhibitors, palbociclib and abemaciclib, were daily administered 7 days after tumor inoculation for 3 weeks, and anti-mouse PD-1 antibody was administered intraperitoneally twice per week (each group n=5). **B, C**. Both abemaciclib and palbociclib showed modest effects for controlling tumor growth. A combination of CDK4/6 inhibitor and PD-1 blockade markedly suppressed tumor growth.

CyTOF displayed that the majority of immune compositions in untreated AB1 tumors were TAMs (**Fig.8**). Treatment of anti-PD-1 antibody led to a modest increase of T cells and a slight decrease of TAMs. Palbociclib treatment led to a significant increase of both B cells and CD4 T cells, and the significant decrease of TAMs. Combination therapy resulted in abundant infiltration of B cells and CD4 T cells and a dramatic decrease of TAMs. Taken together, a combination of CDK4/6 inhibitors (palbociclib or abemaciclib) and a PD-1 inhibitor eradicated anti-PD-1-resistant Cdkn2a(-) AB1 MM tumors with tumor cells through the reprogramming of the tumor-immune microenvironment.

**Figure 8.**
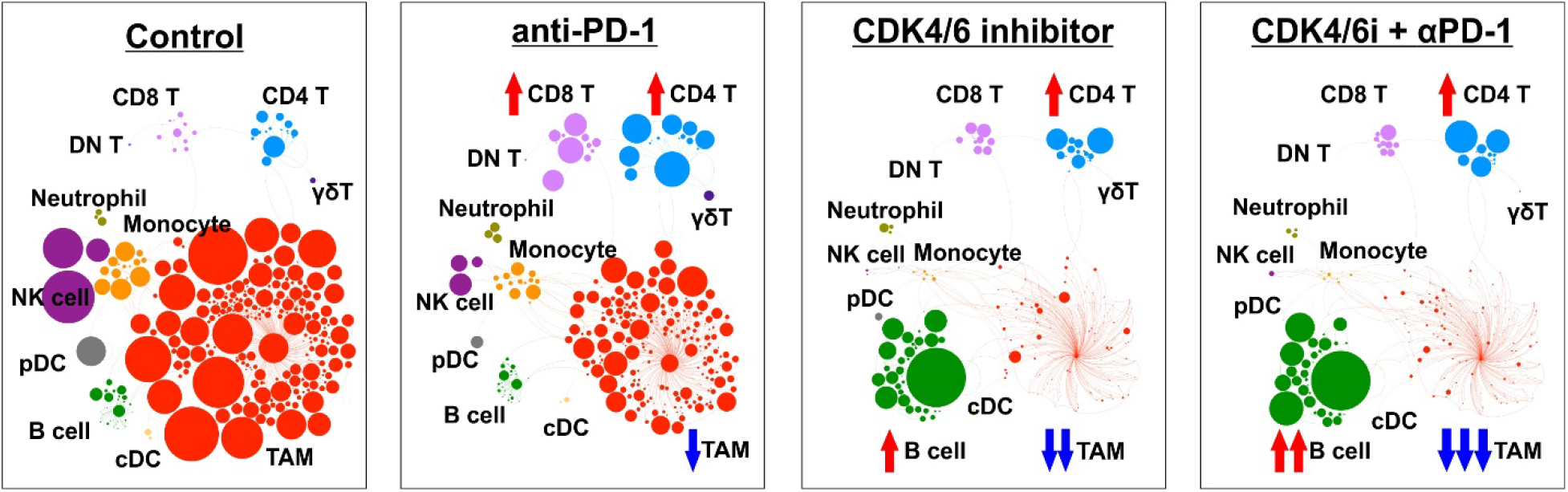
CDK4/6 inhibitors alter the tumor-immune microenvironment. Time of flight mass cytometry showed that most immune compositions in untreated AB1 tumors were tumor-associated macrophages (TAMs). A single treatment of anti-PD-1 led to a modest increase of T cells and a slight decrease of TAMs. Palbociclib treatment led to a significant increase of B cells and CD4 T cells, and a significant decrease of TAMs. Combination therapy resulted in abundant infiltration of B cells and CD4 T cells and a dramatic decrease of TAMs.

## COMMENT

Immune checkpoint inhibitors targeting the PD-1/PD-L1 axis have changed the standard-of-care treatment for a variety of advanced cancers. In MPM, PD-1 inhibitors have recently shown encouraging clinical activity with good tolerability in patients previously treated with chemotherapy.^5–7^ However, the mechanisms of response and resistance to immune checkpoint blockade in human tumors are only beginning to be understood and are still unknown in MPM. Through comprehensive immunogenomic analysis, we uncovered that *CDKN2A* loss is associated with primary resistance to PD-1 blockade. Our *in vivo* experiment using immunocompetent mouse MM model with primary resistance to anti-PD-1 treatment supported that suppressing CDK4/6 activated by *CDKN2A* loss overcomes primary resistance to checkpoint immunotherapy.

Deletions in the 9p21 area containing *CDKN2A* and *CDKN2B* genes are found in 45-52% of MPMs.^20,32,33^ Targeting proliferative cancer cells by CDK4/6 inhibitors may directly eliminate cancer cells, which results in the sensitization of the tumor immune microenvironment to PD-1 blockade by abundant neoantigen exposure. Furthermore, a potential link between CDK4/6 activity and PD-L1 protein stability was recently reported.^34^ In addition to the anti-proliferative effect of CDK4/6 inhibitors on cancer cells, recent studies have indicated that CDK4/6 inhibitors may influence immune cells in the tumor microenvironment, and CDK4/6 inhibitor-induced senescence may enhance the production of a functional chemoattractant for T cells.^35^ A strong relationship between copy number loss of a large region of chromosome 9p and decreased lymphocyte estimates was also reported in melanoma, pancreatic, and head/neck cancers.^36^ The analysis of specimens before and after nivolumab treatment suggests that genetic alterations in *SOX2* and *CDKN2A/CDKN2B* and changes in the tumor microenvironment could be reasons for the acquired resistance to nivolumab observed in the patient with lung cancer.^37^

To face the challenge of primary resistance, constant effort has been made on combination strategies to broaden the responders.^12,38,39^ Furthermore, there has been an extensive search for predictive biomarkers for initial ICI response. PD-L1 expression, tumor mutational burden, tumor-infiltrating lymphocytes, or related gene expression signatures have been explored as potential predictors, and various other markers are currently under investigation.^40^ Our study also provides the potential role of homozygous deletion of *CDKN2A* by fluorescence in situ hybridization as a mechanistic predictive marker to recommend the combination treatment of CDK4/6 inhibitors with ICIs.

Our study limitations include the small number of patients with more advanced disease receiving ICIs to generate the immunologic profile that was applied to the TCGA cohort with earlier disease, requiring further mechanistic investigation with multiple strains to determine the effect of CDK4/6 inhibitors on multiple cellular compositions as well as cancer cells, and uncovering the cause and effect between *CDKN2A* and the alteration of tumor milieu.

## CONCLUSIONS

Through comprehensive immunogenomic approaches, we have identified CDK4/6 as a novel therapeutic target for overcoming primary resistance to PD-1 blockade. These data provide the rationale for undertaking clinical trials of CDK4/6 inhibitors combined with PD-1 blockade in the more than 40% of patients with malignant pleural mesothelioma who demonstrate loss of *CDKN2A*.

## ACKNOWLEDGEMENTS

This work was supported by the Cancer Prevention and Research Institute of Texas (Lee: CPRIT RP200443) and the NIH R37 MERIT Award (Burt: R37CA248478). This project was also supported by the Cytometry and Cell Sorting Core at Baylor College of Medicine with funding from the CPRIT Core Facility Support Award (CPRIT-RP180672), the NIH (CA125123 and RR024574) and the assistance of Joel M. Sederstrom. The authors would like to thank Miriam King, a member of the BCM Michael E. DeBakey Department of Surgery Research Core Team, for editorial assistance during the preparation of this article.

## GLOSSARY OF ABBREVIATIONS

B2M: beta-2-microglobulin
CDKN2A: Cyclin Dependent Kinase Inhibitor 2A
CDK4/6: Cyclin Dependent Kinase 4/6
CR: complete response
CyTOF: time-of-flight mass cytometry
FFPE: formalin-fixed paraffin-embedded
GEO: gene expression omnibus database
ICIs: immune checkpoint inhibitors
IMC: imaging mass cytometry
LOOCV: leave-one-out cross validation
MM: malignant mesothelioma
MPM: malignant pleural mesothelioma
NCBI: the National Center for Biotechnology
PD: progressive disease
PD-1: Programmed Cell Death 1
PD-L1: Programmed Cell Death 1 Ligand 1
PR: partial response
RPKM: reads per kilobase of transcript, per million mapped reads
SD: stable disease
SVM: support vector machine
TCGA: The Cancer Genome Atlas
TAMs: tumor-associated macrophages

