## Supplementary for "Inhibition of CDK4/6 Overcomes Primary Resistance to PD-1 Blockade in Malignant Mesothelioma"

**Table of Contents**


| Section | | Page |
|---|---|---|
| **Supplementary Methods** | | 3 |
| **Supplementary Tables** | | |
| Table S1 | Characteristics of Malignant Pleural Mesothelioma patients receiving PD-1 blockade. | 7 |
| Table S2 | Antibody panel for human and mouse imaging mass cytometry. | 8 |
| Table S3 | Antibody panel for mouse time-of-flight mass cytometry. | 9 |
| Table S4 | Genes for Anti-PD-1 resistant gene signature. | 10 |

### Supplementary methods

#### mRNA Data Analysis

FFPE samples were sliced to 10 μm thickness, and 3-5 slices were put into a 1.5-ml tube. After deparaffinization with xylene, total RNA was extracted with a RecoverAll Total Nucleic Acid Isolation Kit (Ambion, Austin, TX, USA) according to the manufacturer's instructions. RNA quality was assessed with an Agilent 2100 bioanalyzer and the RNA 6000 NanoChip kit (Agilent Technologies, Santa Clara, CA), and RNA quantity was determined using an ND-2000 spectrophotometer (NanoDrop Technologies, Wilmington, DE, USA). To identify the mRNA expression profile of MPM samples, mRNA microarray experiments were performed with a HumanHT-12 v4 Expression Beadchip Kit (Illumina, San Diego, CA, USA). Using a TotalPrep RNA Amplification Kit (Illumina), we labeled and hybridized 750 ng of total RNA according to the manufacturer's protocols. After beadchips were scanned with a BeadArray Reader (Illumina), microarray data were analyzed with the Robust MultiArray Average algorithm and implemented quantile normalization with log2 transformation of gene expression intensities with Biometric Research Branch-Array Tools version 4.5.1[1] and the R script from the Bioconductor project (www.bioconductor.org). Then, we selected the human mRNAs and adjusted data with median values for genes and arrays, respectively. An unsupervised hierarchical clustering algorithm was applied using the uncentered correlation coefficient as the measure of similarity and the method of average linkage (Cluster 3.0).[2] Java Treeview 1.60 (Stanford University School of Medicine, Stanford, CA, USA) was used for tree visualization.[3] Microarray data have been deposited in the National Center for Biotechnology Information's (NCBI) Gene Expression Omnibus (GEO): GSE99070.

#### Imaging mass cytometry (IMC)

The formalin-fixed paraffin-embedded (FFPE) tissues were sectioned at a 5-μm thickness for IMC. FFPE tissues on charged slides were stained with 1:100 diluted antibody cocktails (concentration of each antibody = 0.5mg/mL) recommended by the user's manual. The slides were scanned in the Hyperion Imaging System (Fluidigm). At least four regions of interest in > 4mm$^2$ width were scanned at 200 Hz. All IMC antibodies are listed in **Supplementary Table 2.**

#### Time-of-flight mass cytometry (CyTOF)

***Single cell preparations***

Tumors were finely minced and digested in unsupplemented RPMI 1640 (without glutamate) using a mouse Tumor Dissociation Kit (Miltenyi Biotec Inc., Auburn, CA, USA, Cat# 130-096-730) in 50 mL Falcon tubes for 30 minutes in the 37°C rotating incubator. Cells were then filtered through a 70 μm cell strainer (Corning Life Sciences Plastic, Cat# 07201431), washed, and lysed in ACK lysing buffer (Life Technologies, Cat# A049201). After centrifuging cells with unsupplemented RPMI media for 5 min at 400×g in 20°C and washing two times, supernatants were carefully suctioned off. The cell pellet underwent a final washing with 40 mL supplemented cell culture media (RPMI with FBS), and cells were refiltered through a 70 μm cell strainer. After centrifuging the cells for 5 min at 400×g in 20°C, the supernatant was carefully suctioned off. Some cells were placed in freezing media (FBS with 7% DMSO) and cryopreserved in -80°C freezer storage after cell counting with cell counter (Countess II FL, Life Technologies, Cat# AMQAF1000). For long-term preservation, the cryovials were transferred into a liquid nitrogen tank at -196°C.

***CyTOF***

CyTOF Antibodies were chosen to facilitate the identification of immune cell types, stromal cells, and cancer cells. Antibodies were either purchased preconjugated from Fluidigm (http://maxpar.fluidigm.com/product-catalog-metal.php) or purchased purified and conjugated in-house using MaxPar® X8 Polymer Kits (Fluidigm) according to the manufacturer's instructions by the University of Texas MD Anderson Cancer Center Cytometry Core Facility. Surface and intracellular antibody cocktails were prepared for the staining of all samples for CyTOF. Cryopreserved single cell preparations are stabilized for 6 hours in 37°C incubator after thawing. Five minutes before antibody staining, Cell ID™ -cisplatin (MaxPar®) was added to assess cell viability. Dead cells were stained with Cell-ID Cisplatin according to the manufacturer's protocol. Five hundred thousand cells were resuspended in Maxpar® Cell Staining Buffer (Fluidigm, Cat# 201068) in individual 5 mL tubes for each sample to be barcoded. Mass-tag cellular barcoding using the Cell-ID 20-Plex Pd Barcoding Kit (Fluidigm®, Cat# 201060) was performed. Briefly, $0.5 \times 10^6$ cells from each sample were barcoded with distinct combinations of stable palladium (Pd) isotopes chelated by isothiocyanobenzyl-EDTA in 0.02% saponin in PBS. After washing, cells were resuspended in 1 mL Fix I Buffer (Fluidigm®, Cat# 201065), and incubated for 10 minutes at room temperature (RT). After washing twice with 1 mL of Barcode Perm Buffer (Fluidigm, Cat# 201057), each sample was resuspended to be barcoded completely in 800 µL Barcode Perm Buffer. Barcodes were resuspended completely in 100 µL Barcode Perm Buffer and transferred to the appropriate samples. After mixing the sample immediately and completely, the samples were incubated for 30 minutes at RT. After washing twice with 1 mL of Maxpar® Cell Staining Buffer, the samples were resuspended in 100 µL Maxpar® Cell Staining Buffer, and all barcoded samples were combined into one tube. For the first cohort of 12 MPM patients, cells were washed and incubated with extracellular antibodies for 30 minutes at RT and washed before being fixed and permeabilized in 1× Fix I buffer. The samples were then stained with intracellular antibodies for 30 minutes at RT and washed. Ten minutes before finishing staining with intracellular antibodies, 0.125 nM Cell-ID™ Intercalator-Ir (Fluidigm®, Cat# 201192B) in Maxpar® Fix and Perm Buffer (Fluidigm®, Cat# 201067) was added. Cell-ID™ Intercalator-Ir is a 4 cationic nucleic acid intercalator that contains naturally abundant Iridium (191Ir and 193Ir) and is used for identifying nucleated cells in CyTOF analysis. All CyTOF antibodies are listed in **Supplementary Table 3.**

***CyTOF Data Acquisition***

After washing cells with PBS and MilliQ water, stained cells were analyzed on a mass cytometer (CyTOF3™ mass cytometer, Fluidigm) at an event rate of 400 to 500 cells per second. Data files for each sample were normalized with Normalizer v0.1 MCR and gated. The bead standards were prepared immediately before analysis, and the mixture of beads and cells were filtered through a filter cap FACS tubes before analysis. All mass cytometry files were normalized together using the mass cytometry data normalization algorithm, which used the intensity values of a sliding window of these bead standards to correct for instrument fluctuations over time and between samples. Barcodes were deconvoluted using the Debarcoder® software (Fluidigm®).

***CyTOF Data Analysis***

Total live nucleated cells were used for all analyses and visualized using Spanning-tree Progression Analysis of Density-normalized Events (SPADE)-based SCAFFOLD (Single-Cell Analysis by Fixed Force- and Landmark-Directed) map generation as a mixture of human guided knowledge and automated clustering algorithm.[4-6] To characterize immune cells sorted from all the cells in tumors, all 296 nodes, or clusters of single cells with similar phenotypes, were organized in 11 phenotypes. Eighteen cellular phenotypes were manually defined by a panel of 40 antibodies: B cells (CD45+CD19+B220+), CD4 T cells (CD45+CD3+TCR-β+CD4+), CD8 T cells (CD45+CD3+TCR-β+CD8+), double negative (DN) T cells, (CD45+CD3+TCR-β+CD4-CD8-), γδT cells (CD45+CD3+ TCR-β-TCR-γδ+), monocytes (CD45+CD3-CD11b+CD115+), tumor-associated macrophages (TAMs; CD45+CD3-CD64+F4/80+SiglecF-), plasmacytoid dendritic cells (CD45+CD3-CD19-CD11c+B220+CD317+), conventional DC (CD45+CD3-CD19-CD11c+B220-MHCII+), neutrophils (CD45+CD3-CD19-CD11b+Ly-6G+), and NK cells (CD45+CD3-CD19-CD64-CD49b+).

Based on SPADE results, we generated hierarchical inferences combined with the representative node. A cluster of similar patterns of protein expression was regarded as the same phenotype, and the most fitted node to each phenotype was regarded as the representative node. After assignment of all nodes to each phenotype, nodes were connected and displayed with single cell analyses by fixed force and landmark directed (SCAFFOLD) maps for creating a reference map from high-dimensional single-cell data, facilitating comparisons across samples.[7] SCAFFOLD maps were generated as previously reported.[7] Briefly, a graph is constructed by connecting together the nodes representing the manually gated landmark populations and then connecting to them the nodes representing the cell clusters as well as connecting the clusters to one another. Each node is associated with a vector containing the median marker values of the cells in the cluster (unsupervised nodes) or gated populations (landmark nodes). Edge weights are defined as the cosine similarity between these vectors after comparing the results from the implementation of several distance metrics. Each circle represents a node, a population of cells with the same pattern of expression, and the size of the node represents the percentage of cells in each experiment. The clusters for all the tissues are combined in a single graph with edge weights defined as the cosine similarity between the vectors of median marker values of each cluster. All the pair-wise distances are calculated. The graph is then laid out using the ForceAtlas2 algorithm in Gephi 0.9.1 (https://gephi.org). In this representation of the cellular immune system of the mouse lung, each node is a population of cells with a similar pattern of protein expression, and the size of the node correlates with the average number of cells in the corresponding cell population. To overlay the additional samples on the SCAFFOLD map, the position and identity of the landmark nodes were fixed and the clusters of each sample were connected to the landmark nodes as described above.

**Supplementary Table 1.** Characteristics of Malignant Pleural Mesothelioma patients receiving PD-1 blockade.

| **Variable** | MPM |
|---|---|
| **Number of patients** | 8 |
| **Age**, yr. Median (range) | 62 (38-70) |
| **Gender** | |
| Male | 6 (75 %) |
| Female | 2 (25 %) |
| **Histology** | |
| Epithelial | 4 (50 %) |
| Biphasic | 3 (37.5 %) |
| Sarcomatoid | 1 (12.5 %) |
| **Pre-immunotherapy treatment** | |
| Cisplatin + Pemetrexed | 5 (62.5 %) |
| Carboplatin + Pemetrexed | 3 (37.5%) |
| **Preoperative PD-L1,** % Median (range) | 5 (0-40) |
| **Nivolumab** | 8 (100 %) |
| **Modified RECIST criteria (>6 months)** | |
| CR | 3 (37.5%) |
| PR | 1 (12.4%) |
| PD | 4 (50%) |

**Supplementary Table 2.** Mouse and human antibody panel for imaging mass cytometry.

| Label | Mouse | | | | Human | | | |
|---|---|---|---|---|---|---|---|---|
| | Target | Clone | Source | Cat. No | Target | Clone | Source | Cat. No |
| **141Pr** | **p-Rb (S807/S811)** | J112-906 | BD | 558389 | **CD38** | EPR4106 | DVS-Fluidigm | 3141018D |
| **142Nd** | **Caspase 3, cleaved** | D3E9 | DVS-Fluidigm | 3142004A | **CD19** | 6OMP31 | DVS-Fluidigm | 3142014D |
| **143Nd** | **Vimentin** | RV202 | DVS-Fluidigm | 3143029D | **Vimentin** | D21H3 | DVS-Fluidigm | 3143027D |
| **144Nd** | **TGFB1** | 310-390 aa, C-terminal | LifeSpan BioSciences. | LS-B14345 | **TGFB1** | 310-390 aa, C-terminal | LifeSpan BioSciences. | LS-B14346 |
| **145Nd** | **CD68** | FA-11 | BioLegend | 137002 | **CTLA-4** | CHO cells | Selleckchem | A2001 |
| **146Nd** | | | | | **CD16** | EPR16784 | DVS-Fluidigm | 3146020D |
| **147Sm** | **F4/80** | A3-1 | AbD Serotec | MCA497GA | **CD163** | EDHu-1 | DVS-Fluidigm | 3147021D |
| **148Nd** | **Pan-Cytokeratin** | C11 | DVS-Fluidigm | 3148020D | **Pan-Keratin** | C11 | DVS-Fluidigm | 3148020D |
| **149Sm** | **CD11b** | EPR1344 | DVS-Fluidigm | 3149028D | **CD11b** | EPR1344 | DVS-Fluidigm | 3149028D |
| **150Nd** | **PD-L1** | 10F.9G2 | Biolegend | 50164172 | **IFNgamma** | B27 | Biolegend | 10761-890 |
| **151Eu** | **IgM** | RMM-1 | DVS-Fluidigm | 3151006B | **CD31** | EPR3094 | DVS-Fluidigm | 3151025D |
| **152Sm** | **p-H2AX, p-gamma-H2AX** | N1-431 | BD | 560443 | **CD45** | CD45-2B11 | DVS-Fluidigm | 3152016D |
| **153Eu** | **CD44** | IM7 | DVS-Fluidigm | 3153029D | **CD44** | IM7 | DVS-Fluidigm | 3153029D |
| **154Sm** | **p21, WAF1/Cip1** | CP74 | Sigma | P1484 | **CD11c** | Polyclonal | DVS-Fluidigm | 3154025D |
| **155Gd** | **FOXP3** | polyclonal | LifeSpan BioSciences. | LS-C357445 | **FoxP3** | 236A/E7 | DVS-Fluidigm | 3155016D |
| **156Gd** | **CD4** | polyclonal | LifeSpan BioSciences. | LS-C359266 | **CD4** | EPR6855 | DVS-Fluidigm | 3156033D |
| **158Gd** | **E-Cadherin** | 2.4E+11 | DVS-Fluidigm | 3158029D | **E-Cadherin** | 24E10 | DVS-Fluidigm | 3158029D |
| **159Tb** | **p-AKT** | M89-61 | BD | 560397 | **CD68** | KP1 | DVS-Fluidigm | 3159035D |
| **160Gd** | **p-PI3K (p85/p55)** | Poly | CST | 4228BF | **Vista** | D1L2G | DVS-Fluidigm | 3160025D |
| **161Dy** | **CD19** | polyclonal | LifeSpan BioSciences. | LS-B13077 | **CD20** | H1 | DVS-Fluidigm | 3161029D |
| **162Dy** | **CD8A** | Polyclonal | LifeSpan BioSciences. | LS-C662748 | **CD8a** | C8/144B | DVS-Fluidigm | 3162034D |
| **163Dy** | **c-Myc** | D84C12 | CST | 5605BF | **PD-L1** | CHO cells | Selleckchem | conjugated |
| **164Dy** | **Arginase-1** | D4E3M | DVS-Fluidigm | 3164027D | **C-Myc p67** | 9E10 | DVS-Fluidigm | 3164025D |
| **165Ho** | **IFNg** | XMG1.2 | DVS-Fluidigm | 3165003B | **PD-1** | CHO cells | Selleckchem | conjugated |
| **166Er** | **IL-4** | 11B11 | DVS-Fluidigm | 3166003B | **CD45RA** | HI100 | DVS-Fluidigm | 3166028D |
| **167Er** | **c-Myc** | D84C12 | CST | 5605BF | **Granzyme B** | EPR20129-217 | DVS-Fluidigm | 3167021D |
| **168Er** | **Ki67** | B56 | DVS-Fluidigm | 3168022D | **Ki-67** | B56 | DVS-Fluidigm | 3168022D |
| **169Er** | **IL-10** | Polyclonal | LifeSpan BioSciences. | LS-B7432 | **IL-10** | Polyclonal | LifeSpan BioSciences. | LS-B4913-100 |
| **170Er** | **CD3E** | polyclonal | LifeSpan BioSciences. | LS-C343957 | **CD3** | Polyclonal, C-Terminal | DVS-Fluidigm | 3170019D |
| **171Yb** | **PD-1** | 29F.1A12 | Bioxcell | BP0273 | **CD27** | EPR8569 | DVS-Fluidigm | 3171024D |
| **172 Yb** | | | | | **CX3CR1** | 2A9-1 | DVS-Fluidigm | 3172017B |
| **173 Yb** | **CD80** | RM80 | LifeSpan BioSciences. | LS-C44886 | **CD45RO** | UCHL1 | DVS-Fluidigm | 3173016D |
| **174Yb** | **PTEN** | 17.A | Invitrogen | PIMA512278 | **HLA-DR** | YE2/36 HLK | DVS-Fluidigm | 3174023D |
| **175Lu** | | | | | **CD86** | BU63 | Abcam | ab213044 |
| **176Yb** | **Lyz** | EPR2994(2) | Abcam | ab185129 | **Lyz** | EPR2994(2) | Abcam | ab185129 |

**Supplementary Table 3.** Mouse antibody panels for time-of-flight mass cytometry.

| Label | Surface | | | |
|---|---|---|---|---|
| | Target | clone | Source | Cat. No |
| 89Y | CD45(Ms) | 30-F11 | DVS-Fluidigm | 3089005B |
| 115In | CD4(Ms) | RM4-5 | BioLegend | 100506 |
| 127IdU | S-phase cell cycle | | | |
| 139La | CD45.2 | 104 | BioLegend | 109843 |
| 141Pr | Ly-6G | 1A8 | DVS-Fluidigm | 3141008B |
| 142Nd | CD11c | Polyclonal | DVS-Fluidigm | 3142003B |
| 143Nd | CCR2 | 475301 | R&D | MAB55381 |
| 144Nd | CD115 | AFS98 | DVS-Fluidigm | 3144012B |
| 145Nd | CD69 | H1.2F3 | DVS-Fluidigm | 3145005B |
| 146Nd | CD8a (Ms) | 53-6.7 | DVS-Fluidigm | 3146003B |
| 147Sm | CD62L | MEL-14 | BioLegend | 104443 |
| 148Nd | CD11b (Ms) | M1/70 | DVS-Fluidigm | 3148003B |
| 149Sm | CD19 (Ms) | 6D5 | DVS-Fluidigm | 3149002B |
| 150Nd | CD25 | 3C7 | DVS-Fluidigm | 3150002B |
| 151Eu | CD64 | X54-5/7.1 | DVS-Fluidigm | 3151012B |
| 152Sm | CD3e | 145-2C11 | DVS-Fluidigm | 3152004B |
| 153Eu | CD274, PD-L1 | 10F.9G2 | DVS-Fluidigm | 3153016B |
| 154Sm | Cytokeratin(pan) | C-11 | BioLegend | 628602 |
| 155Gd | CD278, ICOS | C398.4A | BioLegend | 313502 |
| 156Gd | Vimentin | RV202 | DVS-Fluidigm | 3156023A |
| 158Gd | Foxp3 | FJK-16s | DVS-Fluidigm | 3158003A |
| 159Tb | CD279, PD-1 | 29F.1A12 | DVS-Fluidigm | 3159024B |
| 160Gd | TCRgd | GL3 | eBioscience | 14-5711-82 |
| 161Dy | CD40 | HM40-3 | DVS-Fluidigm | 3161020B |
| 162Dy | Ly-6C | HK1.4 | DVS-Fluidigm | 3162014B |
| 163Dy | CD152, CTLA-4 | 9H10 | BioLegend | 106202 |
| 164Dy | CX3CR1 | SA011F11 | DVS-Fluidigm | 3164023B |
| 165Ho | CD317, PDCA-1 | 927 | BioLegend | 127002 |
| 166Er | CD44 | IM7 | BioLegend | 103002 |
| 167Er | c-Myc | D84C12 | CST | 5605BF |
| 168Er | Ki67 | B56 | BD | 556003 |
| 169Tm | TCRbeta | H57-597 | DVS-Fluidigm | 3169002B |
| 170Er | CD49b, Integrin α2 | HMα2 | DVS-Fluidigm | 3170008B |
| 171Yb | CD80 | 16-10A10 | DVS-Fluidigm | 3171008B |
| 172Yb | CD86 | GL1 | DVS-Fluidigm | 3172016B |
| 173Yb | F4/80 | BM8 | BioLegend | 123102 |
| 174Yb | CD326 (Ms) | G8.8 | BioLegend | 118201 |
| 175Lu | CD38 | 90 | DVS-Fluidigm | 3175014B |
| 176Yb | B220, CD45R | RA3-6B2 | BioLegend | 103202 |
| 195Pt | Cell viability | | | |
| 209Bi | I-A/I-E, MHC-II | M5/114.15.2 | DVS-Fluidigm | 3209006B |

**Supplementary Table 4.** Genes for anti-PD-1 resistance signature.

| # | Gene symbol | P-value | Fold change (log2) | # | Gene symbol | P-value | Fold change (log2) |
|---|---|---|---|---|---|---|---|
| 1 | A1CF | 0.001926017 | -1.760698572 | 80 | MGC16121 | 0.00756817 | -1.613353038 |
| 2 | ABL2 | 0.005778448 | 1.333253888 | 81 | MUC5B | 0.008544592 | 1.142758193 |
| 3 | ACADM | 0.002468075 | -1.726467137 | 82 | MUM1L1 | 0.006036535 | 1.526545127 |
| 4 | ACBD5 | 0.007178588 | 1.248070266 | 83 | MUS81 | 0.00388895 | -1.742211687 |
| 5 | ADAM6 | 0.001893221 | -1.834722179 | 84 | MX1 | 0.004426645 | -1.453808943 |
| 6 | ADAMTS19 | 0.004475869 | 1.686879384 | 85 | MYO7A | 0.000303608 | 1.851147905 |
| 7 | ADRA1A | 0.005377473 | 1.580804029 | 86 | NBPF10 | 0.00885333 | 1.627612367 |
| 8 | AIM2 | 0.002310882 | -1.858689042 | 87 | NEUROG3 | 0.00607667 | -1.769533968 |
| 9 | ALDH1A2 | 0.007933211 | -1.688858537 | 88 | NFATC2IP | 0.002312654 | -1.68286809 |
| 10 | ARGFXP2 | 0.004345199 | -1.729656818 | 89 | NFATC3 | 0.001590321 | 1.784144289 |
| 11 | BBS7 | 0.006448138 | 1.790803639 | 90 | NOL9 | 0.005270846 | 1.688052195 |
| 12 | BCL7A | 0.003063777 | 1.874429641 | 91 | NOTCH2NL | 0.006364424 | 1.673785308 |
| 13 | BEND4 | 0.004932073 | -1.661526942 | 92 | NUP54 | 0.001403774 | 1.738998508 |
| 14 | BRCA1 | 0.009562223 | -1.145010704 | 93 | OPRD1 | 0.004332597 | -0.781215631 |
| 15 | C16orf42 | 0.000197024 | -1.582592059 | 94 | OR10A7 | 0.003273514 | -1.87472641 |
| 16 | C1orf127 | 0.00355508 | -1.254142722 | 95 | OR2A2 | 0.003984551 | 1.76912989 |
| 17 | CDC6 | 0.009232455 | 1.617865007 | 96 | OR6C65 | 0.002509217 | -1.569785762 |
| 18 | CEACAM20 | 0.006254206 | 1.686429388 | 97 | PANK2 | 0.003087108 | -1.796296116 |
| 19 | CLDN6 | 0.006538255 | -1.291318325 | 98 | PCBP2 | 0.003637452 | 1.702620618 |
| 20 | CNRIP1 | 0.008961682 | 1.778101031 | 99 | PCGF2 | 0.009542674 | 1.452956756 |
| 21 | COG3 | 0.006451243 | 1.803291288 | 100 | PCNA | 0.002972182 | 1.846747128 |
| 22 | COL9A1 | 0.005722518 | -1.617914118 | 101 | PGM3 | 0.006890908 | 1.664639855 |
| 23 | CRISP2 | 0.003835624 | -1.447575061 | 102 | PHF12 | 0.008226558 | 1.496812962 |
| 24 | CST4 | 0.001589578 | 1.366581633 | 103 | PNPO | 0.002177928 | -1.896611723 |
| 25 | CYFIP1 | 0.001372974 | 1.850185935 | 104 | PPP2R2B | 2.31528E-05 | 2.015234651 |
| 26 | DBN1 | 0.000386029 | 1.704212311 | 105 | PTK2B | 0.006597816 | -1.729999651 |
| 27 | DEFA5 | 0.008386785 | -1.211668569 | 106 | RAB43 | 0.003885702 | -1.809824223 |
| 28 | DNAH17 | 0.005607152 | -1.464095745 | 107 | RARRES1 | 0.003935857 | -1.614961121 |
| 29 | DNAJA3 | 0.008609746 | -1.7616982 | 108 | RBMY3AP | 0.002218685 | 1.731480297 |
| 30 | EBAG9 | 0.008616486 | -1.687761246 | 109 | RIMBP3C | 0.007058752 | 1.786474302 |
| 31 | EIF5 | 0.003967313 | -1.536055051 | 110 | RNF39 | 0.006316426 | 0.56780024 |
| 32 | ENAH | 0.009775625 | 1.626404245 | 111 | RPS6KA5 | 0.00876094 | -1.473703717 |
| 33 | EPB41 | 0.005642849 | 1.358590261 | 112 | RTN3 | 4.07388E-05 | -1.684739112 |
| 34 | ERAS | 0.00047968 | -1.861129863 | 113 | SCNN1A | 0.006542277 | -1.245391428 |
| 35 | EYA4 | 0.002490164 | -1.532018618 | 114 | SCP2 | 0.001485885 | -1.600593211 |
| 36 | FAM71E2 | 0.001453467 | -1.768660384 | 115 | SERPINB12 | 0.004367128 | 1.826590025 |
| 37 | FBXO24 | 0.001154684 | -1.727137536 | 116 | SIRT5 | 0.007170287 | -1.523277771 |
| 38 | FLJ14107 | 0.0074351 | 1.575618262 | 117 | SKA2 | 0.003611884 | -1.659612421 |
| 39 | FLJ25328 | 0.002024152 | 1.347527517 | 118 | SLC37A3 | 0.008096465 | 1.639160339 |
| 40 | GAB2 | 0.008077743 | -1.66290649 | 119 | SLC4A8 | 0.002591918 | -1.457279481 |
| 41 | GABRG2 | 0.003301371 | -1.596806546 | 120 | SLC6A14 | 0.004968129 | -1.547661562 |

| | | | |
|---|---|---|---|
| 42 | GAS1 | 0.002334935 | 1.815942178 |
| 43 | GCET2 | 0.001721565 | -1.920599081 |
| 44 | GLE1 | 0.009775182 | 1.47130706 |
| 45 | GPM6B | 0.005833483 | 1.794201554 |
| 46 | GPR55 | 0.005585241 | -1.349744764 |
| 47 | H6PD | 0.008329606 | 1.557431585 |
| 48 | HIST1H2BM | 0.005062682 | 1.68876799 |
| 49 | HIVEP3 | 0.005513168 | -1.470160141 |
| 50 | HPS4 | 0.009293869 | -1.655441066 |
| 51 | HSPA1L | 0.005354921 | -1.800870381 |
| 52 | IGLL1 | 0.002762734 | -1.727313285 |
| 53 | IL22RA2 | 0.005159646 | 1.766686677 |
| 54 | IL28A | 0.003103524 | -1.810579399 |
| 55 | IL33 | 0.002354446 | 1.493344196 |
| 56 | INPP4B | 0.000701043 | -1.454378385 |
| 57 | IP6K2 | 0.007251889 | 1.440431504 |
| 58 | IQCA1 | 0.000364482 | 1.803086712 |
| 59 | IRF7 | 0.005788122 | -1.649308796 |
| 60 | ITGA11 | 0.009793455 | 1.550256145 |
| 61 | KCNF1 | 0.00733989 | 1.508769074 |
| 62 | KCNMB2 | 0.003413733 | 1.348292525 |
| 63 | KCNQ2 | 0.002604519 | -1.899825014 |
| 64 | KIAA0319L | 0.007254819 | -1.713436396 |
| 65 | KIF16B | 7.50913E-05 | 2.036376982 |
| 66 | KLF11 | 0.004597693 | 1.583933769 |
| 67 | KLHL6 | 0.004536732 | 1.74601748 |
| 68 | KLRG2 | 0.00971813 | -1.347708588 |
| 69 | LDHA | 0.006152084 | -1.72956763 |
| 70 | LRRC1 | 0.004125048 | -1.827036608 |
| 71 | LRRC34 | 0.002050979 | 1.497367526 |
| 72 | M6PR | 0.005673107 | -1.623539694 |
| 73 | MAN1A2 | 0.003284545 | -1.844261328 |
| 74 | MAPK1IP1L | 0.002582006 | -1.807536403 |
| 75 | MAX | 0.0099241 | -1.707248148 |
| 76 | MBP | 0.002586935 | -1.803133409 |
| 77 | MCRS1 | 0.002048342 | 1.162421057 |
| 78 | METTL7B | 0.006035269 | -1.667458541 |
| 79 | MEX3B | 0.008433225 | 1.631341381 |
| 121 | SLCO1A2 | 0.008498208 | 1.639442258 |
| 122 | SNCB | 0.004915308 | 1.760880392 |
| 123 | SNORA5A | 0.009219678 | -1.741713837 |
| 124 | SNORD49A | 0.002132703 | 1.396865827 |
| 125 | SNORD69 | 0.002274865 | 1.81096929 |
| 126 | SNX20 | 0.001129844 | 1.083066757 |
| 127 | SP110 | 0.003811201 | -1.624202989 |
| 128 | SP140 | 0.004311164 | -1.482442321 |
| 129 | SPATA7 | 0.008893923 | 1.747944822 |
| 130 | SPOCD1 | 0.007706922 | -1.523008143 |
| 131 | SPTBN4 | 0.009027349 | -1.201780935 |
| 132 | SRMS | 0.006774626 | -1.446907584 |
| 133 | ST3GAL6 | 0.00154254 | -1.474764453 |
| 134 | STOML1 | 0.006918576 | -1.589023659 |
| 135 | STXBP1 | 0.003932145 | 1.575588331 |
| 136 | TAAR6 | 0.007863254 | -0.661371267 |
| 137 | TAC4 | 0.006791879 | -1.512292307 |
| 138 | TBPL2 | 0.005093235 | -1.34782936 |
| 139 | TCP11L2 | 0.000416908 | 1.969953914 |
| 140 | TEX9 | 0.006979143 | 1.804585709 |
| 141 | TGFB2 | 0.003352966 | 1.69758668 |
| 142 | TGM5 | 0.006334409 | 1.812920903 |
| 143 | TIAM2 | 0.009361282 | -1.512486564 |
| 144 | TINAGL1 | 0.00273972 | 1.824394479 |
| 145 | TMEM140 | 0.003795543 | -1.596553115 |
| 146 | TMEM170A | 0.006249256 | -1.655719188 |
| 147 | TRIM17 | 0.00870819 | 1.623783169 |
| 148 | TRPV1 | 0.000210797 | -1.985760569 |
| 149 | UBE3C | 0.008646868 | -1.718827377 |
| 150 | UBQLN1 | 0.006427152 | 1.137157331 |
| 151 | USP17 | 0.005235872 | -1.27888599 |
| 152 | VCL | 1.68493E-05 | 1.919701923 |
| 153 | WIPF2 | 0.002724628 | -1.789207042 |
| 154 | WNT1 | 0.005650741 | 1.574307092 |
| 155 | WNT5B | 0.00546917 | -0.935186586 |
| 156 | WWOX | 0.000635214 | 1.683374 |
| 157 | ZNF45 | 0.007844801 | 1.450532206 |
| 158 | ZNF74 | 0.003772907 | -1.644150598 |
